# A natural disaster exacerbates and redistributes disease risk across free-ranging macaques by altering social structure

**DOI:** 10.1101/2023.07.17.549341

**Authors:** Alba Motes-Rodrigo, Gregory F. Albery, Josue E. Negron-Del Valle, Daniel Philips, Cayo Biobank Research Unit, Michael L. Platt, Lauren J.N. Brent, Camille Testard

**Author notes:** Corresponding author **Corresponding author contact**: Camille Testard; Address: 16 Divinity Ave, Cambridge, MA, 02138, USA. These authors contributed equally to this work (Alba Motes-Rodrigo, Gregory F. Albery). **Statement of authorship:** CT, AMR and GFA conceptualised the study; JN and DP collected the behavioural data used in this study; CT, AMR and GFA analysed the data; MLP, LJNB secured funding for this work; AMR, GFA and CT drafted the manuscript with input from all authors. All authors read and approved the manuscript. **Data and code accessibility statement:** Data and code is available on GitHub at https://github.com/camilletestard/Cayo-Maria-Disease-Modeling.

## Abstract

Climate change is intensifying extreme weather events, with severe implications for ecosystem dynamics. A key behavioural mechanism whereby animals may cope with such events is by increasing social cohesion to improve access to scarce resources like refuges, which in turn could exacerbate epidemic risk due to increased close contact. However, how and to what extent natural disasters affect disease risk via changes in sociality remains unexplored in animal populations. By modelling disease spread in free-living rhesus macaque groups (*Macaca mulatta*) before and after a hurricane, we demonstrate doubled pathogen transmission rates up to five years following the disaster, equivalent to an increase in pathogen infectivity from 10% to 20%. Moreover, the hurricane redistributed the risk of infection across the population, decreasing status-related differences found in pre-hurricane years. These findings demonstrate that natural disasters can exacerbate and homogenise epidemic risk in an animal population via changes in sociality. These observations provide unexpected further mechanisms by which extreme weather events can threaten wildlife health, population viability, and spillover to humans.

## MAIN TEXT

Natural disasters have devastating consequences for both man-made and natural ecosystems, leading to casualties, displacement, and psychological distress^1,2^. In group-living animals, including humans, social relationships help individuals cope with such challenges^3^ and altering social patterns can be a critical behavioural mechanism by which animals adapt to disaster-induced ecosystem degradation^2,4–6^. For instance, human social groups become more transitive^7^ and cohesive^8^ following natural disasters, while dolphin groups become more modular^9^ and primate social networks more densely connected^6^. Changes in social structure can also be caused by increased mortality following disasters^10,11^ or through behavioural adjustments to new resource landscapes and ecological challenges^6^. Regardless of the cause, changes in the topography of social relationships among group members following natural disasters can have important consequences for a range of downstream processes like selection, population viability, and disease dynamics, none of which are well-understood in the context of disasters^12^. This knowledge gap severely reduces our ability to anticipate the consequences and mechanisms of global change for human and non-human social species.

Social structure plays a fundamental role in governing the spread of contagious entities such as innovations^13^, intestinal microbes^14^, and pathogens^15^. In particular, epidemic risk strongly depends on the population’s demographics and connectedness as well as between-individual variation in social interactions, among other factors^16^. Previous studies have shown that perturbations that alter a group’s social structure^15^ can slow the spread of infections in wild animals^17,18^, or speed up pathogen spread if the group becomes more densely connected^19^. Natural disasters constitute large ecological perturbations and given that climate change is predicted to increase the frequency and severity of extreme weather events^20^, it is imperative that we evaluate the long-term effects of such disasters on epidemic risk in both animal and human populations. Better understanding this process could help identify a route by which global change is threatening animal conservation^21^ as well as influencing the risk of pathogen spillover into humans^22^.

Natural disasters often concentrate displaced individuals at high densities in limited safe spaces known as refugia^12,23–25^, which then drives greater contact rates – a process that has been highlighted as a cause of infectious disease outbreaks in humans and potentially other animals^26,27^. However, disasters’ effects on social structure have yet to be explicitly linked with changes in epidemic risk in animals. This knowledge-gap exists partly because disasters are unpredictable such that researchers are rarely able to *plan* to study them^12,28^. Furthermore, to build this link between natural disasters, social structure and epidemic risk, individual-level social data is required. However, almost all known studies of disasters’ demographic and behavioural effects have occurred at the population level^29,30^, which do not permit quantifying social relationships across the disaster event. Individual-level data is fundamental in this regard because the costs of disasters are unlikely to fall evenly across a population, and will affect some individuals more than others, at least partly due to their social environment^31^. For example, disaster-induced epidemics could act more heavily on low-status individuals who might have a compromised immune system due to lack of access to high-quality foods^32,33^. Alternatively higher-ranking individuals with access to more social partners might be exposed to a higher risk of disaster-induced epidemics^31,34^. Age classes might also differ in susceptibility to disaster-induced disease risk due to immunosuppression and social selectivity in older individuals^35,36^. Different social environments or strategies based on sex could also lead to variation in disease risk due to different rates of exposure to pathogens^37^. Moreover, the distribution of contacts across a population can have strong and nonlinear effects on epidemic risk – i.e., by influencing superspreader events^38^.

Here, leveraging a long-term individual-based dataset from a natural experiment, we address this gap using epidemiological simulations applied to empirical social interaction data from hundreds of free-ranging rhesus macaques (*Macaca mulatta*) collected from 5 years before until 5 years after a natural disaster, Hurricane Maria (**Figure S1** & **Table S1**). Specifically, we quantify (i) how changes in social structure following Hurricane Maria influence individual risk of pathogen acquisition, and (ii) how individual-level sociodemographic factors mediate infection risk before and after the hurricane. This study therefore takes full advantage of the power of long-term individually-based studies in disease ecology^39,40^, and presents a unique individually-based data set of an animal population affected by a natural disaster.

## METHODS

### 1. Study system

We studied a semi-free-ranging population of rhesus macaques living on Cayo Santiago Island, Puerto Rico (1809 N, 65 44 W). Rhesus macaques live in multi-male multi-female groups with a polygynandrous mating system. Male rhesus macaques disperse from their natal group around sexual maturity, but females typically remain in their natal group for their entire lives, such that social groups are stably composed of matrilines^41,42^. Rhesus macaques are notable for their high frequency and severity of aggression and absence of reconciliation between conspecifics when compared to other macaque species, which leads to a social organisation that can be described as both highly “despotic and nepotistic”^43,44^.

The population of rhesus macaques used for this study has been continuously monitored since it was established in 1938 following the release of 409 animals originally captured in India. Cayo Santiago is managed by the Caribbean Primate Research Center (CPRC), which supplies food to the population daily and water *ad libitum*. There is no contraceptive use and no medical intervention aside from tetanus inoculation when animals are weaned yearlings. Animals are free to aggregate into social units as they would in the wild and there are no natural predators on the island.

On September 20^th^ 2017, Hurricane Maria made landfall on Cayo Santiago. This hurricane was one of the deadliest and most destructive natural disasters in Caribbean history. Within a few days, over 60% of Cayo Santiago’s vegetation was destroyed^6^ and five years later (Spring 2023) most of it still has not recovered^17^. Severe deforestation greatly reduced shade availability, a fundamental resource for behavioural thermoregulation in the intense Caribbean heat^6^. As a result, monkeys concentrated in fewer, smaller shady areas as well as in exposed areas thus increasing the frequency of close-proximity contacts^17^ (**Figure S2**).

Subjects for this study were adult males and females (at least six years old), individually recognizable by tattoos, ear notches, and facial features. We used seven social groups, TT, F, KK, V, R, HH and S, for which there was behavioural data either before and/or after Hurricane Maria, between 2013 and 2022. Our dataset included 790 unique adult individuals. We used multiple available years of observational data (HH: 2014, 2016; R: 2015, 2016; F: 2013, 2014, 2015, 2016, 2017; KK: 2013, 2015, 2017, 2018; V: 2015, 2016, 2017) to characterise social networks before the hurricane (‘‘pre-hurricane’’). Post-hurricane, our sample included observational data from 2018, 2019, 2021 and 2022 (S: 2019; TT: 2022; F: 2021, 2022; V: 2018, 2019, 2021, 2022; KK: 2018). See **Figure S1** and **Table S1** for details on sample size within each group and year.

### 2. Behavioural data collection

In all years of data collection except 2018 (year immediately following Hurricane Maria) and three social groups in 2016, 2017 and 2019 (HH, KK and S respectively), behavioural data were collected with 10-min focal animal samples. At the 0-, 5-, and 10-min marks of the focal follow, we collected instantaneous scan samples during which we recorded the identity of all adults within two metres (i.e., in proximity). For groups HH in 2016, KK in 2017 and S in 2019, we collected 5-min focal animal samples with instantaneous scan samples at the beginning and end of the focal follow, the data collection protocol was otherwise identical to other non-hurricane years. We balanced data collection on individuals across time of day and across months to account for temporal variation in behaviours. Data-collection hours were from 6am to 2:30pm due to CPRC operating hours. To collect behavioural data before 2020 we used Teklogic Psion WorkAbout Pro handheld computers with Noldus Observer software. After 2020, we used the Animal Observer App on Apple iPads.

In the year following Hurricane Maria (from November 2017 to September 2018), damage resulting in inconsistent access to electricity in Puerto Rico imposed the adoption of a downgraded means of recording data using basic tablets. We recorded group-wide instantaneous scan samples at 10-min intervals. For all animals in view of an observer, we recorded the identity of all adults within two meters (i.e., in proximity)–similarly to instantaneous scans recorded during focal follows prior to the hurricane. Observers were given 15 mins to complete a group-wide scan session, were required to stand a minimum of 4 m from monkeys and, because of very good visibility of these terrestrial animals, were able to identify them at distances upward of 30 m.

Our subjects were observed over a mean (SD) of 2.61 (1.70) years, and did not have to be observed both before and after the hurricane to be included in the analysis. In all years except 2018, we included on average 84.32 (36.96) scan observations embedded within focal follows per individual per year. In the year post-hurricane (November 2017 - September 2018), we collected 533.36 (127.41) scans per individual and no focal follows. Note that because of storms and Hurricane Maria, data collection stopped on August 31, 2017 and didn’t resume until November 2nd, 2017.

Dominance ranks for individuals were determined separately for each group and year. Rank was also determined separately for males and females. For males, the direction and outcome of win-loss agonistic interactions recorded during focal animal samples or during ad libitum observations of a given year was used to determine rank for that year. For females, rank was determined using both outcomes of win-loss agonistic interactions and matriline rank. Female macaques inherit their rank from their mothers, and female ranks are linear and relatively stable over time^41^. In order to account for group size, dominance rank was defined by the percentage of same sex individuals outranked, and ranged between 0 to 100 (0 = lowest rank, outranks 0% of same sex individuals; 100 = highest rank, outranks 100%). We classified animals as either ‘high’, “medium” or ‘low’ ranking based on the percentage outranked scale. Monkeys were classified as “high” ranking if they outranked > 80% of the monkeys of their group/sex, “medium” ranking if they outranked 50-80% of their group and “low” ranking if they outranked < 50% of monkeys of their group/sex.

### 3. Bayesian estimating of social networks using BISoN

For each group and year separately, we estimated social networks based on proximity interactions captured during scan samples.

Networks are usually constructed by taking a normalized measure of sociality, such as the proportion of sampling periods each pair spends engaged in a social behavior (e.g. the Simple Ratio Index; SRI). These normalized measures ensure that pairs that are observed for longer are not erroneously determined to be more social^45^. This is important because observing social events can be challenging and uniform sampling over all pairs is not always possible^46^. Though normalized measures of sociality will not be biased by sampling time, they will be accompanied by varying levels of certainty. For example, edge weights will be treated as a certain 0.5 for both a case where individuals have been seen together once and apart once, and equally where individuals have been together 100 times and apart 100 times. Despite there being considerably more certainty around the values of the latter example than the former, methods to estimate and propagate uncertainty through social network analyses remain largely unused.

In this study, we generated weighted Bayesian networks using the BISoN framework and bisonR package^47^. This framework allowed us to account for uncertainty in the edges connecting individuals in the network based on how often they were sampled and, more importantly, propagate this uncertainty to subsequent analyses. Proximity networks were estimated based on data collected through scan sampling^48^ across all groups and years. Edges in the networks represented the number of times a pair of individuals were observed in proximity relative to the total observation effort for the dyad (i.e., total scans individual A + total scans individual B), or the probability of interaction. Given that the observed interaction data was binary (i.e. for every observation sample, was the focal animal in proximity to another monkey? Y/N) we modelled the uncertainty around the edges using as prior a beta distribution with alpha = 0.1 and beta = 0.1. Proximity networks were undirected.

### 4. Pathogen spread simulation

To quantify how hurricane-induced behavioural changes influenced infectious disease risk, we simulated epidemics of a hypothetical pathogen through the annual networks of each macaque group. This approach has been previously deemed the most effective to forecast how pathogens might spread through a population and impact wildlife health^21^, and has been used in a range of contexts to investigate the epidemiological consequences of perturbations (e.g. in ants^18^). Here, we used 1000 Bayesian networks (BISoN package^47^) of each group-by-year combination as simulation replicates in order to account for the uncertainty of dyadic weight estimates in the observed proximity networks. For each simulation we generated a new pathogen infectivity (P)_I_, drawn randomly from a normal distribution (Mean=0.15, SD=0.04, range=0.000001-0.3). This distribution captures a wide range of airborne pathogen transmission risks, from very low to quite high (up to 30% probability of infection). In each simulation, we chose a random individual at timestep 0 to be infected with the hypothetical pathogen. As such, there were three sources of random variation at the initiation of each simulation: the structure of the network (drawn from the Bayesian model posterior), the first infected individual, and the infectivity of the pathogen. In each simulation timestep, each infected individual transmitted the pathogen to its contacts with probability A*B, where A equals the probability that two individuals interact (i.e., the weight of the dyadic connection in the network) and B equals a binomial coin flip with probability P_I_. Each simulation continued until either all individuals were infected or 10,000 timesteps elapsed and revealed the mean time step at which individuals in each group-year replicate became infected.

We fitted a linear mixed model (LMM) to our simulations’ outputs to determine the size and significance of the hurricane’s effect. For this model, each data point was a simulation result (N=24000, comprising 1000 simulations across each of 24 group-by-year combinations) and the response variable was the mean timestep at which individuals in that replicate were infected. The explanatory variables included in the model were: i) the P_I_ value of the pathogen in the simulation; ii) a binary factor denoting whether the simulation included a network from before or after Hurricane Maria and iii) the random intercept of the group-by-year combination. This model formulation was conservative in its ability to detect a hurricane effect, as the group:year random effect and the hurricane effect were confounded, such that some of the variation accounted for by the former was likely originated from the latter. To put the hurricane effect’s magnitude in context, we compared its estimate from the linear model with the estimate for the P_I_ value’s effect (Figure 2C). All analyses were carried out in R (Version 4.2.3).

### 5. Modelling inter-individual differences in epidemic risk

Next, we tested whether inter-individual differences in epidemic risk amongst the population changed pre-to-post hurricane, with faster infection in the population representing a higher epidemic risk^21^. We fitted a gaussian linear mixed model to our simulations’ outputs, mean timestep at which individuals were infected (i.e., a measure of infection speed), to determine the effect of individual characteristics–namely social status, age and sex–on pathogen infectivity before and after the hurricane. Fixed effects included hurricane status (pre or post), age (z-transformed to a mean of 0 and standard deviation of 1), sex (Male or Female) and social status (categorised as Low, Medium and High rank), as well as the interactions between hurricane status and individual characteristics. We also included group nested within individual ID as random effects to account for repeated measures and between-group differences. Pairwise differences between rank levels were calculated using the function emmeans from the package of the same name^49^.

To examine whether the hurricane shuffled the order of individuals’ connectedness in the population, we examined individual-level repeatability of connectedness across the study period. To do so, we followed a protocol used in red deer to identify between-individual variation in seasonality^50^. We quantified repeatability by fitting individuals’ connection network strength as a response variable with Year, Population, and Pre-Post Maria as fixed explanatory variables. We included individual identity as a random effect, and then quantified the proportion of variance accounted for by this random effect (i.e., repeatability; we label this ID Overall Model). Additionally, we added an ID:PrePost interaction as a random effect, which allowed us to identify whether the same individuals with strong connections pre-hurricane had strong connections post-hurricane (ID:Hurricane Model). A large value for the variance accounted for by this random effect would demonstrate a strong interaction effect – i.e., that individuals shifted substantially in the rank order of connection strength before *versus* after the hurricane. Finally, to confirm repeatability of connection strengths within each time period (before or after) we split the data into these two datasets, and then re-ran the repeatability analysis (ID Before and ID After Models).

### 6. Quantifying inter-individual differences in proximity network strength and degree pre- and post-hurricane

To further investigate the source of inter-individual differences in disease risk between ranks pre- and post-hurricane, we quantified individuals’ proximity network strength, or the propensity to be in close proximity to other individuals, using the ‘strength’ function in igraph on BISoN-generated proximity networks (Methods section 3). This function calculates the sum of weights of an individual’s connections in the network. We also quantified individuals’ proximity network degree, or the number of proximity partners, using the ‘degree’ function in igraph^51^.

To quantitatively assess inter-individual differences in proximity strength and degree pre- and post-hurricane, we ran two separate linear models for individual network strength and degree. In both models the response variable was standardised and we included hurricane status, age, sex and social status (categorised as low, medium and high) as main fixed effects, as well as the interactions between hurricane status and each socio-demographic parameter. This way we could test whether inter-individual differences in degree or strength changed pre-to-post hurricane. We also included individuals nested within social groups as random intercepts to account for repeated measures and group differences. To quantify changes in status-related differences in degree and strength pre-to-post hurricane we estimated marginal means using the ‘emmeans’ function in R. We contrasted low, medium and high ranking individuals pre- and post-hurricane.

For all models we ran the following assumption checks to ensure model estimates were accurately estimated using the function ‘check_model’ from R package performance^52^: posterior predictive checks, homogeneity of variance, colinearity of fixed effects, normality of residuals, normality of random effects. We did not find violations of the assumptions checked.

## RESULTS

### Speed of disease spread doubles after hurricane Maria

Following Hurricane Maria individuals became infected on average 1.93 times faster (i.e., in roughly half the number of simulation time steps) than before the storm (95% CI 3.05-1.20; P=0.002, **Figure 1A**) due to their greater social connectedness to conspecifics. Faster infection was detected each year following the hurricane compared to pre-hurricane years (**Figure 1B**) and at equivalent pathogen infectivities, individual infection was faster post- than pre-hurricane (**Figure 1C**). The hurricane’s effect on the speed of pathogen transmission was equivalent to a 10% absolute increase in pathogen infectivity from 0.1 to 0.2, which led to 1.97-2.01 times faster infection spread (*P*<0.001; **Figure 1D**). These simulations reveal substantially higher epidemic risk after the hurricane, implying social responses to natural disasters can strongly exacerbate population vulnerability to infectious disease outbreaks.

**Figure 1.**
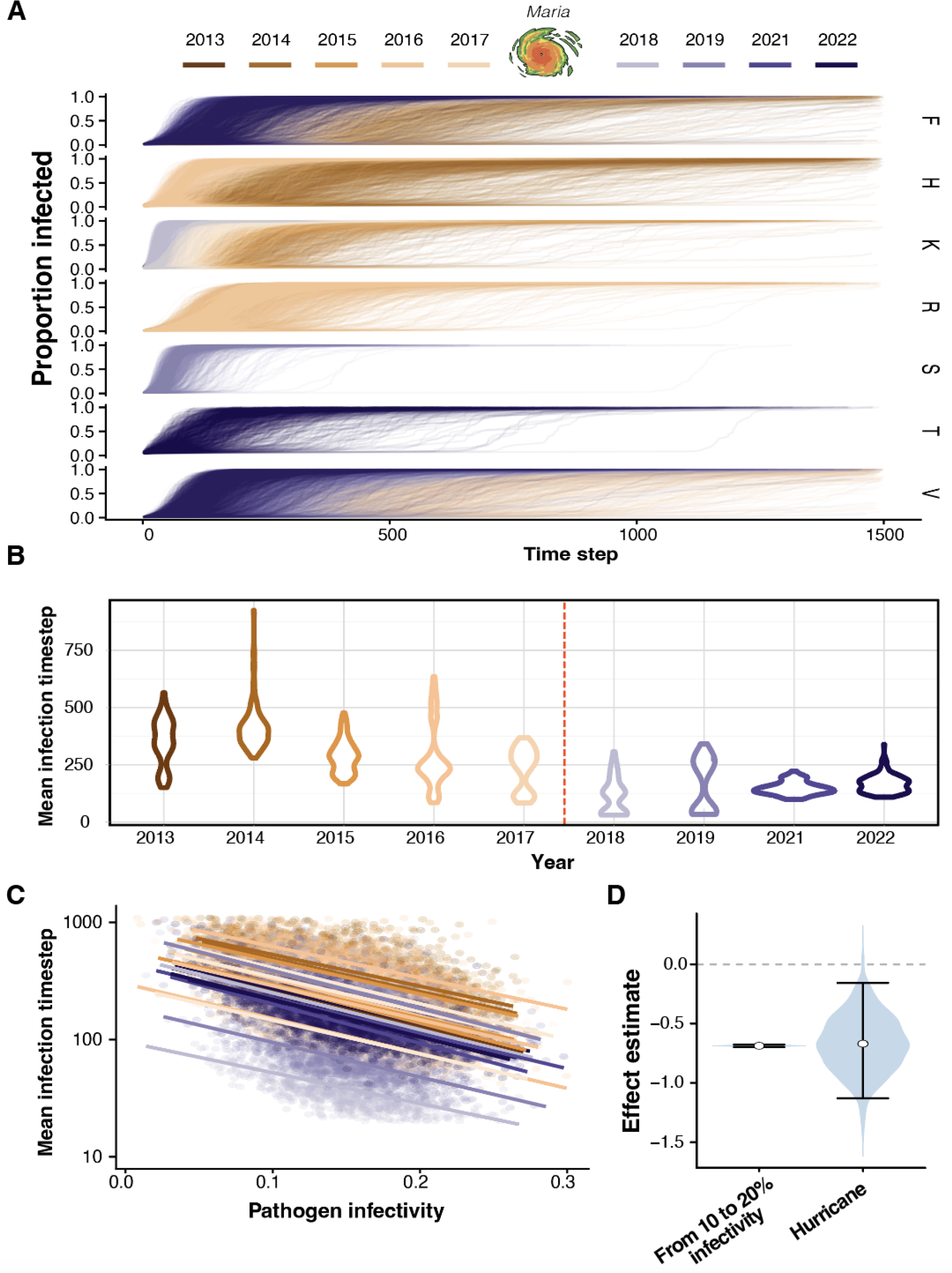
**Simulated epidemic risk as a function of year, pathogen infectivity, social group and individual characteristics**. Simulation outputs are grouped into year and group replicates. Each year is represented by a different colour, with oranges indicating pre-hurricane years and purples post-hurricane years. (**A**) Each line represents one of 1000 simulations carried out on each group-year replicate (N=24,000), depicting the proportion of infected individuals at each simulation timestep. Post-hurricane years (purples) reached higher proportions of infected individuals markedly faster than pre-hurricane years (oranges). Each macaque social group is a different row and the group’s name is on the right of each row. (**B**) Distribution of mean individual infection timesteps separated by years. (**C**) Each point represents the mean simulation time step at which individuals become infected in a given simulation (N=24,000) as a function of pathogen infectivity. Solid lines represent models fitted to each group-by-year replicate, coloured by year as in Panel A. Pathogen infectivity was associated with substantially earlier mean infection timestep on a log_10_-scale. (**D**) The effect estimate for a 10% increase in infectivity is given next to the estimate for the effect of the hurricane. The blue violins represent the probability distribution of the posterior effect estimate; the error bars the 95% credibility interval and the points the mean.

### Disease risk is homogenised after Maria

We found that inter-individual differences in disease susceptibility related to social status were reduced after the hurricane. While low ranking individuals were on average 54.51 steps slower to get infected than high ranking individuals in the years pre-hurricane (contrast estimate between high and low-ranking individuals pre-hurricane = -54.0, SE = 5.53, *p*<0.001, Figure 2A) this status-related difference nearly halved after the hurricane (contrast estimate between high and low-ranking individuals post-hurricane = -25.4, SE = 8.47, *p*=0.033, Figure 2A). Furthermore, the pre-hurricane differences of middle-ranking individuals with high and low-ranking individuals in terms of mean infection time step disappeared after the hurricane (Table S2). Age did not affect epidemic risk before or after the hurricane and sex differences were only mildly attenuated after the hurricane (not significant, Figure 2A).

**Figure 2.**
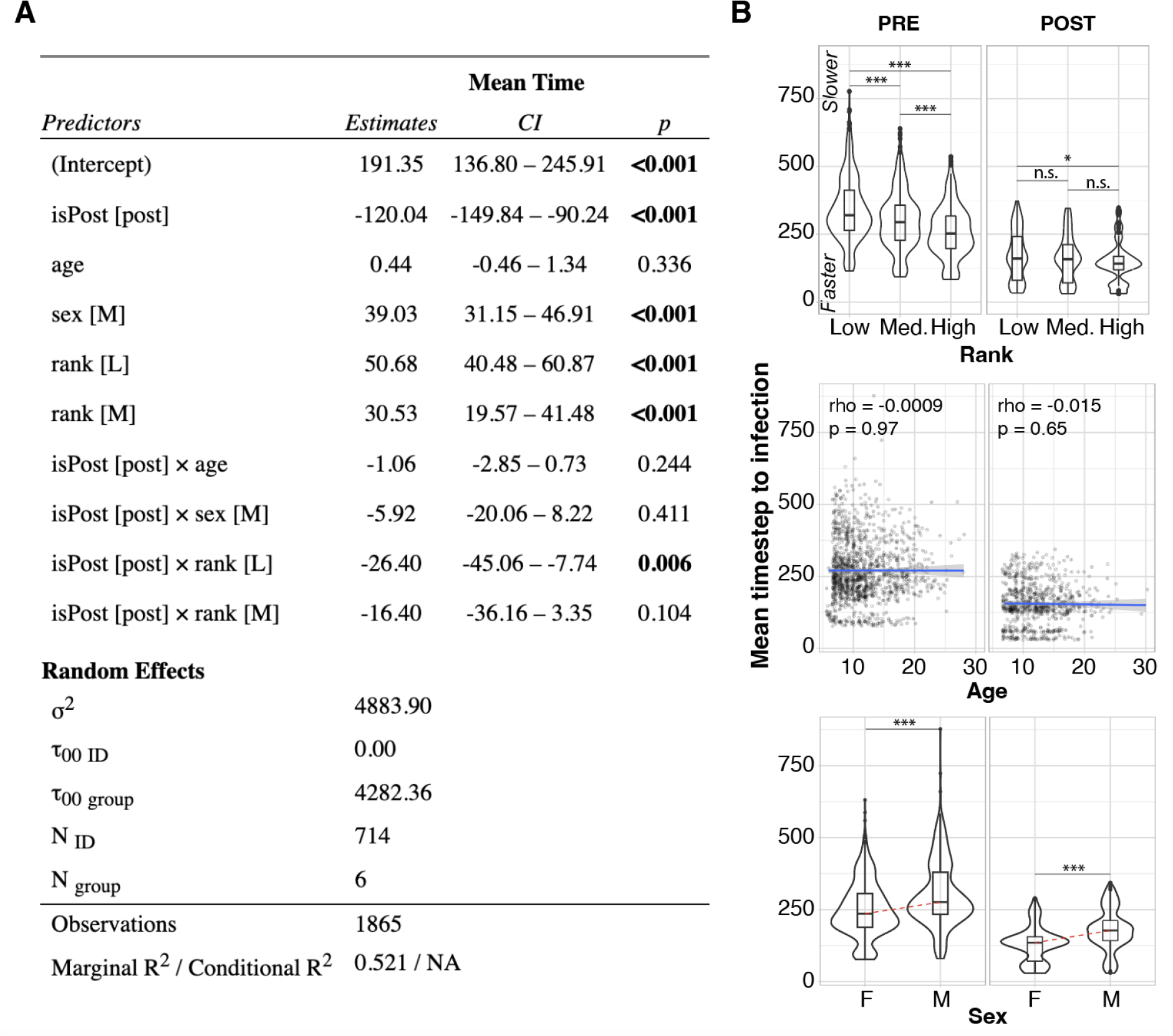
**Hurricane Maria levels the playing field in terms of speed of infection among individuals of different ranks**. A). Gaussian linear mixed model output table testing the effect of individual socio-demographic factors on exposure to disease (“Mean Time”) before vs. after the hurricane (see **Methods**). B) Mean simulated timestep to infection across social status (top), age (middle) and sex (bottom). Statistical significance determined based on post-hoc marginal means on linear mixed model. Rho: pearson correlation coefficient; ***: *P*<0.01; *: *P*<0.05; n.s.: not significant; Med: “Medium:; pre: pre-hurricane; post: post-hurricane, L: low-ranking, M in panel A: middle-ranking; M in panel B: males; F: females; NID: number of individuals; Ngroup: number of groups; σ:mean random effect variance.

Next, we investigated which network properties explained the attenuation of social status-related differences in disease exposure post-hurricane. Although differences in proximity strength between low and high-ranking individuals after the hurricane were exacerbated after the hurricane (contrast estimate between high and low-ranking individuals pre-hurricane: 0.31, SE=0.07; post-hurricane: 0.11, SE=0.10; Figure 3), we found a lower difference in the number of proximity partners between these groups after the disaster (contrast estimate between high and low-ranking individuals pre-hurricane: 0.41, SE=0.075; post-hurricane: 0.76, SE=0.10; Figure 3).

**Figure 3.**
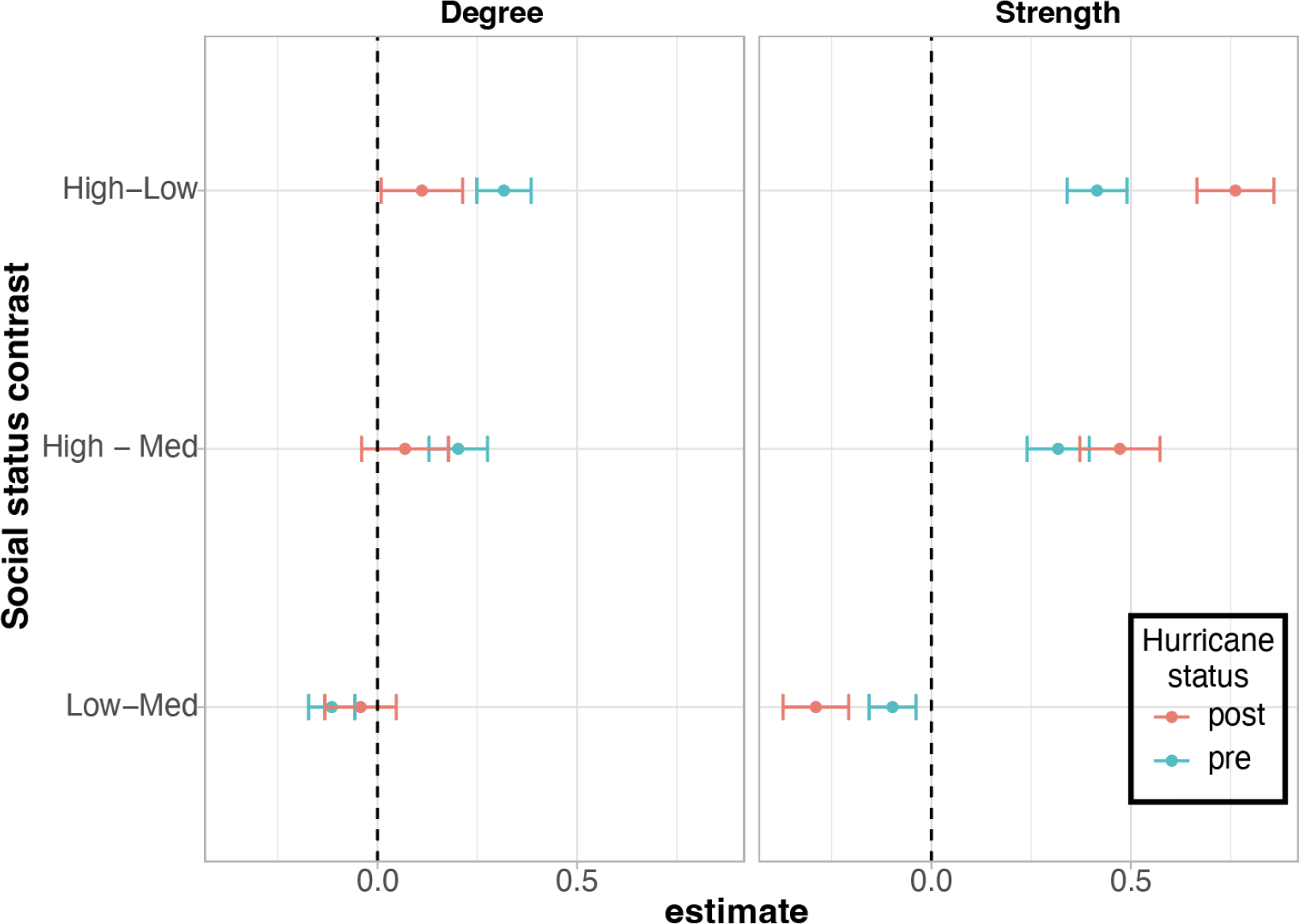
Levelling of disease risk across the macaque social hierarchy is explained by a closer number of proximity partners. Marginal means from linear mixed models of proximity network degree and strength pre- and post-hurricane. Pre-hurricane (pre): blue; Post-hurricane (post): red.

To further explore how the hurricane altered the distribution of relative connectedness and epidemic risk across the population, we quantified between-individual repeatability of behaviour; see Figure S3. We found that individuals’ network connection strengths were repeatable across the study period (R=0.16; 0.09-0.21), but that fitting an ID:hurricane interaction effectively removed this repeatability (R=0.002, 0.00-0.03). The variance accounted for by the ID:hurricane interaction itself was large (R=0.76; range 0.72-0.79), indicating that individuals’ rank order of network strength greatly changed after the hurricane compared to before. That is, the same individuals were not necessarily well-connected before *versus* after the hurricane relative to the rest of the population. Supporting this observation, the repeatability for the period before and after the hurricane were much higher than the overall period (R=0.41 and 0.53 respectively), demonstrating that the hurricane’s shuffling of connection strengths had a profound impact on the distribution of social connectedness across the population (Figure S3).

## DISCUSSION

Our study demonstrates that natural disasters can increase the magnitude and alter the distribution of epidemic risk in a free-ranging animal population. Specifically, the macaques of Cayo Santiago doubled their risk of infection due to higher proximity following Hurricane Maria. Furthermore, the social changes that took place following the hurricane homogenised the epidemic risk of individuals within groups, reducing previous variation in infection risk caused by social status.

This study represents the first quantification of how natural disasters impact epidemic risk via changes in social structure in a closed system where migration is not possible. Similar closed systems are found in island-dwelling species, extreme dietary specialists with geographical restriction (ecological islands) and non-migratory species. As such, our results may generalise to other species similarly dependent on geographically-restricted environmental resources potentially affected by natural disasters.

While the impact of natural disasters on human health are relatively well-understood^53,54^, their influence on the resilience of animal populations remains understudied. By applying epidemiological simulations to empirically-derived social networks pre- and post-hurricane, we provide a first demonstration of how natural disasters can alter infection disease risk via changes in social structure. Our results imply that, if a pathogen invades a population after a natural disaster and reduces host survival or reproduction, it could disproportionately decrease its viability, posing an important conservation threat in addition to the short-term, direct effects of the natural disaster (i.e., increased mortality).

Changes in social structure following Maria was an adaptive behavioural response to altered resource availability^55^ (i.e. shade). The impact this response had on disease risk in our study population illustrates how natural disasters can jeopardise the survival of animal populations indirectly up to five years following the event. These findings demonstrate the importance of considering the long-term consequences of natural disasters, and of habitat disturbance more generally, for the health and survival of free-ranging animals.

The hurricane reduced social inequalities in individual infection times, highlighting that natural disasters can impose surprising levelling effects on epidemic risk. This homogenization of disease susceptibility across social status was due to changes in the number of social partners, not in strength of relationships with those partners. Whereas high-ranking individuals spend more time close to select partners after the hurricane, low-ranking individuals adopted a strategy of “get close to more individuals” and almost matched high-ranking individuals in terms of number of proximity partners. This strategy for low-ranking individuals made them more exposed to disease than they were before the hurricane. In humans, vulnerability of low socio-economic status individuals to disasters have generally been related to lack of access to resources such as space, clean water and food^56^. Here, we describe another mechanism by which low-status individuals show a disproportionate vulnerability to disasters: differences in social strategies to cope with the consequences of the event.

In addition to impacting population viability, greater pathogen prevalence among wildlife in the aftermath of a natural disaster could increase the rate at which humans encounter infected animals or pathogens in the environment. This, in turn, increases the probability of zoonotic spillover^57^, which has become progressively more frequent in the last century^58^. The threat of spillover could be further exacerbated following natural disasters due to infrastructure damage – e.g. to water systems^59^– and increased population density in safe areas^58^. Similarly, if the epidemic erodes social structure and causes animals to behave differently or disperse, the probability of zoonotic spillover may further increase, posing an important public health risk^23^. Taken together, these findings expand our understanding of the role of natural disasters in governing ecosystem health, and imply that future plans for conservation and public health should incorporate disasters’ effects on wildlife health emerging from behavioural changes.

## Acknowledgements

We thank the Caribbean Primate Research Center staff for their important roles in data collection and in the maintenance of Cayo Santiago.

## SUPPLEMENTARY INFORMATION

**Figure S1.**
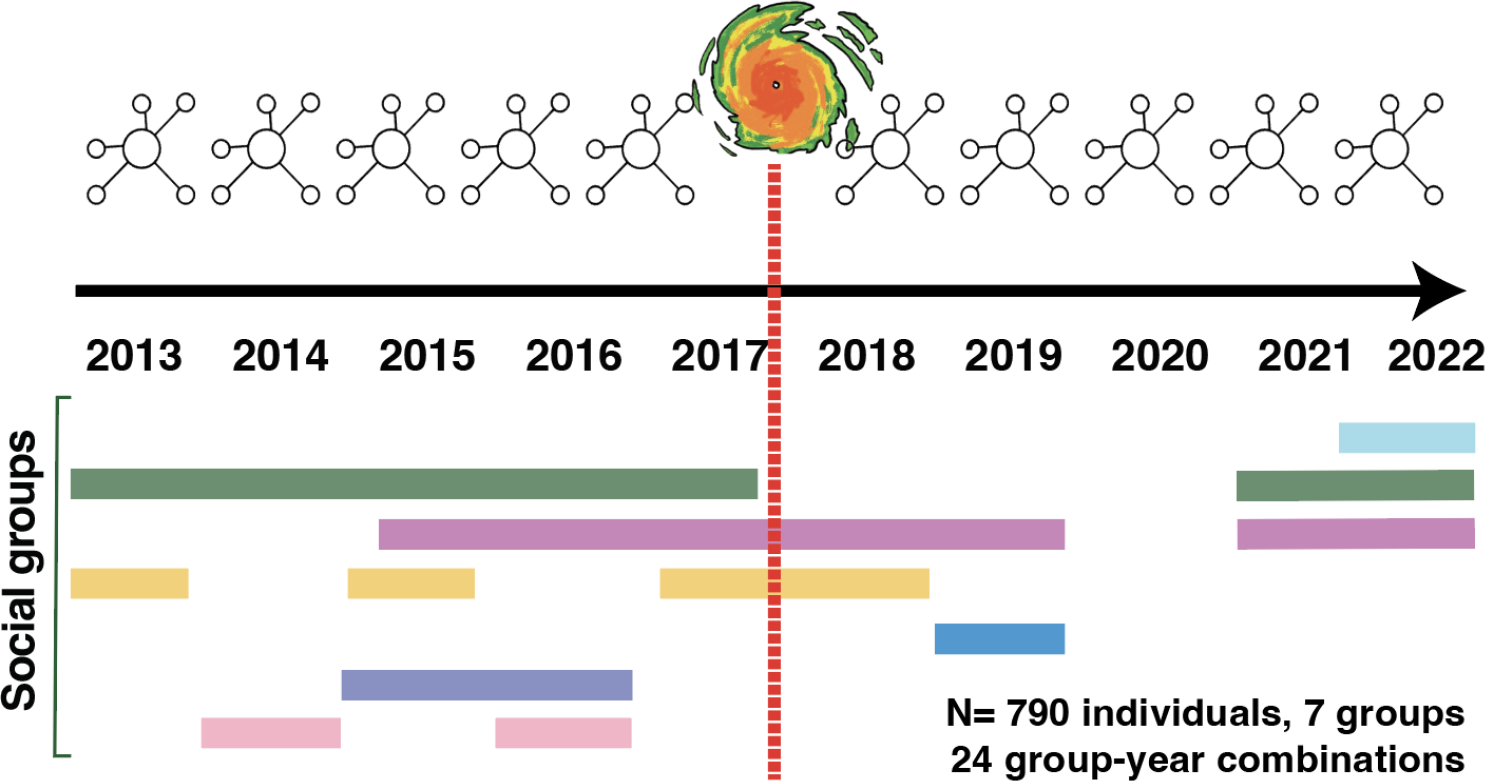
Coverage of data collection across social groups and years. Adapted from *Testard et al. 2023*.

**Figure S2.**
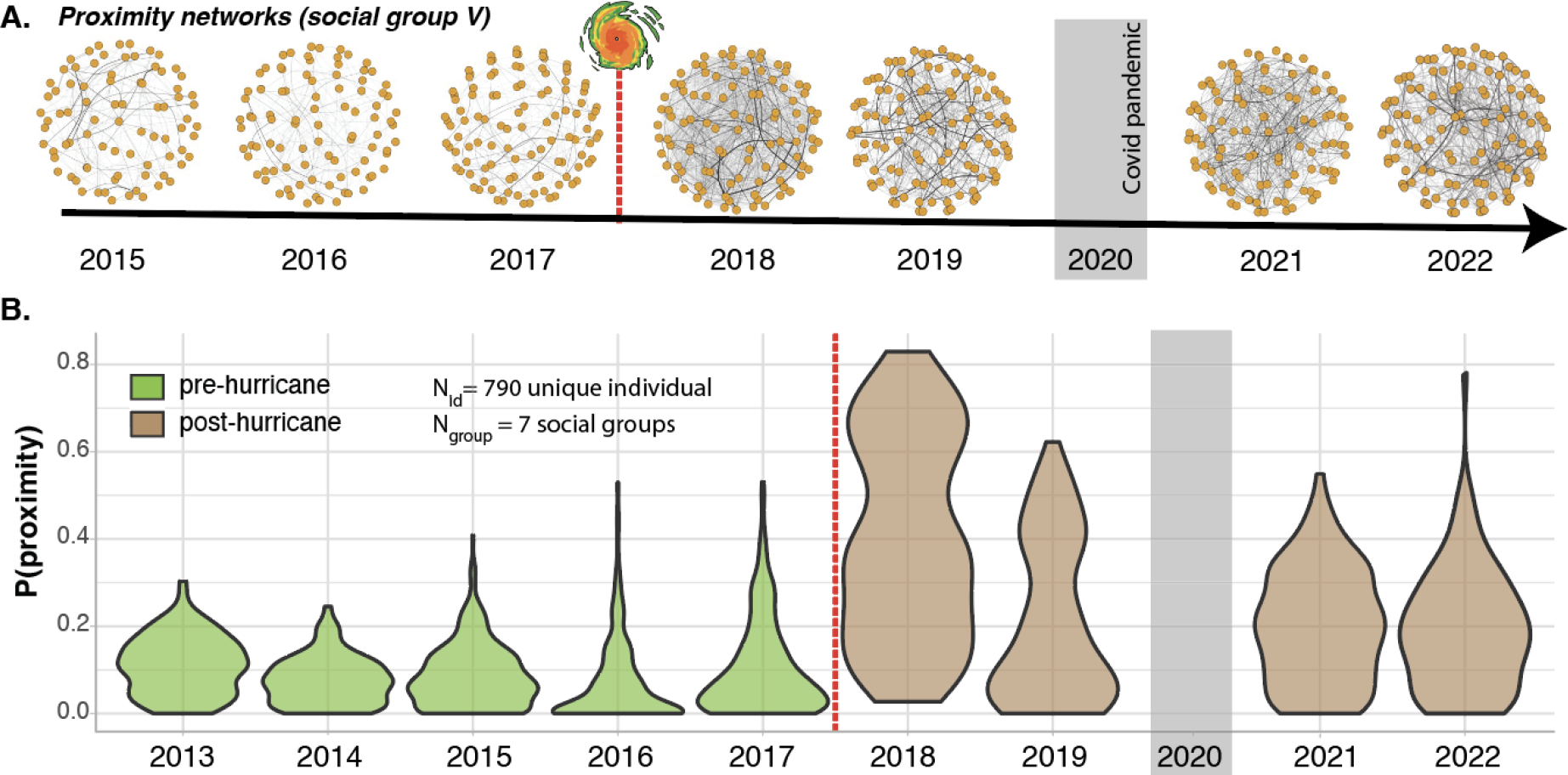
Cayo Santiago macaques exhibit a persistent increase in social tolerance after Hurricane Maria. (**A**) Proximity networks for social group V from 2013 to 2022, excluding 2020 due to the COVID19 crisis are shown as example networks. Each node represents an individual and edges represent the probability for two individuals to be in proximity to one another. The thicker the line the higher the probability of proximity. Red dotted line marks Hurricane Maria. (**B**) Probability of being observed in proximity to another monkey from 2013 to 2022, across 790 individuals in 6 social groups, 24 group-year combinations. Adapted from *Testard et al. 2023*

**Figure S3.**
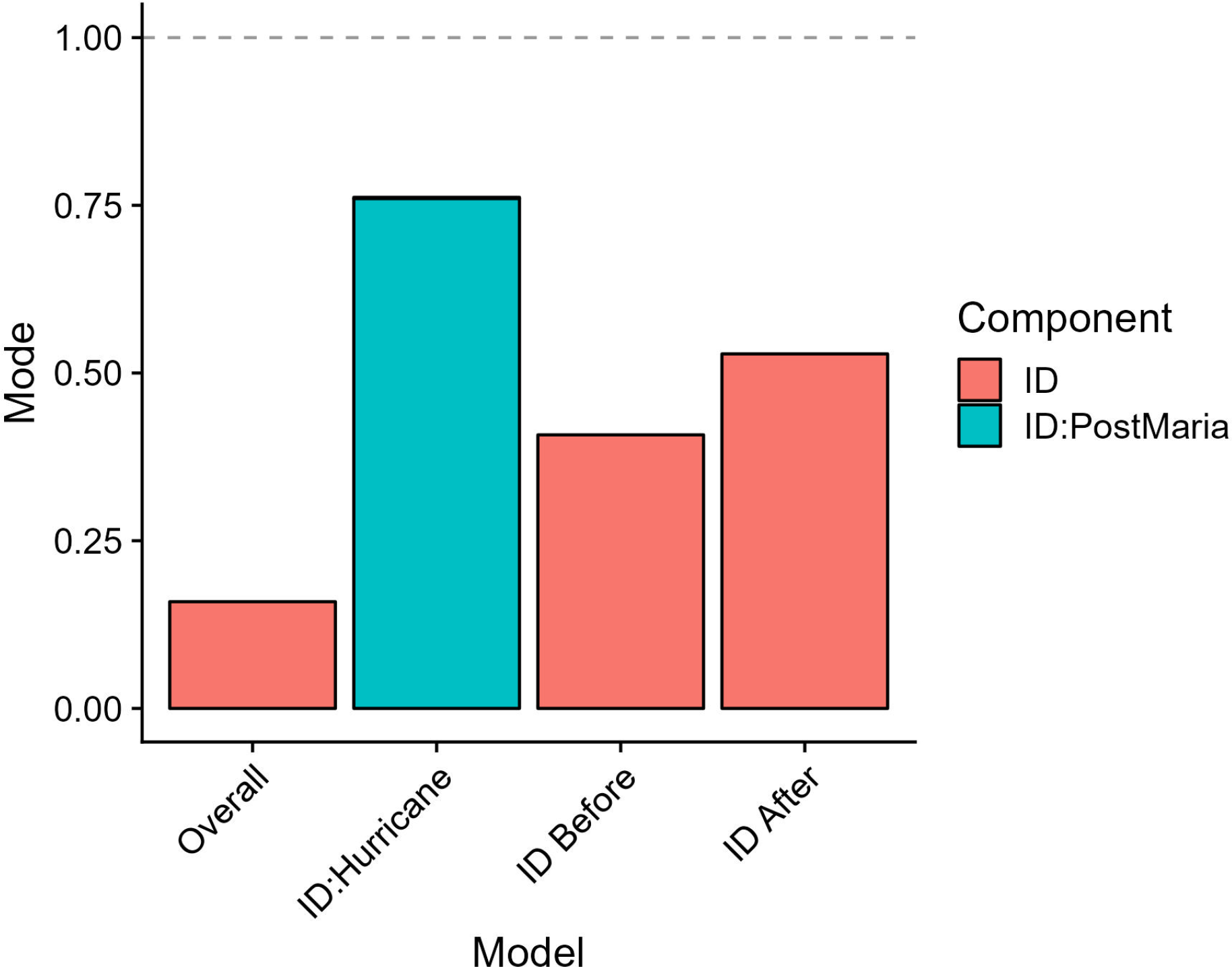
Repeatability of connection strength across the hurricane. Repeatability was relatively high overall (first column), but was much higher when isolated to only examine the data collected before or after the hurricane (third and fourth columns). The extremely large effect of the ID:Hurricane interaction (second column) denotes that individuals’ connection strengths changed substantially between pre- and post-hurricane, essentially removing overall repeatability from the model.

**Table S1.** Sample sizes by group and year. Note that an individual can be sampled in multiple years.

|  | 2013 | 2014 | 2015 | 2016 | 2017 | 2018 |
| --- | --- | --- | --- | --- | --- | --- |
| F | 100 | 106 | 103 | 123 | 113 | 0 |
| HH | 0 | 45 | 0 | 69 | 0 | 0 |
| KK | 30 | 0 | 54 | 0 | 61 | 63 |
| R | 0 | 0 | 94 | 125 | 0 | 0 |
| S | 0 | 0 | 0 | 0 | 0 | 0 |
| TT | 0 | 0 | 0 | 0 | 0 | 0 |
| V | 0 | 0 | 80 | 90 | 102 | 67 |

**Table S2.** Contrasts between rank categories in terms of mean infection timesteps before and after the hurricane. P values are corrected for multiple testing using the Tukey correction. SE = standard error, df = degrees of freedom.

| Contrast | estimate | SE | df | t.ratio | P. value |
| --- | --- | --- | --- | --- | --- |
| pre H - post H | 145.91 | 8.64 | 1851.57 | 16.88 | 1.84E-11 |
| pre H - pre L | -53.97 | 5.53 | 831.52 | -9.76 | 0.00 |
| pre H - post L | 120.55 | 7.05 | 1405.21 | 17.11 | 0.00 |
| pre H - pre M | -31.83 | 5.94 | 953.22 | -5.36 | 1.57E-06 |
| pre H - post M | 131.17 | 7.66 | 1551.71 | 17.13 | 0.00 |
| post H - pre L | -199.88 | 7.87 | 1705.99 | -25.39 | 2.85E-12 |
| post H - post L | -25.36 | 8.47 | 1683.55 | -2.99 | 0.03 |
| post H - pre M | -177.74 | 8.18 | 1751.48 | -21.72 | 5.85E-12 |
| Post H - post M | -14.74 | 8.91 | 1764.37 | -1.65 | 0.56 |
| pre L - post L | 174.52 | 6.08 | 1852.65 | 28.72 | 1.78E-11 |
| pre L - pre M | 22.14 | 4.72 | 1165.66 | 4.69 | 4.56E-05 |
| pre L - post M | 185.14 | 6.78 | 1829.40 | 27.33 | 1.49E-11 |
| post L - pre M | -152.38 | 6.47 | 1663.51 | -23.54 | 1.35E-12 |
| post L - post M | 10.62 | 7.31 | 1785.65 | 1.45 | 0.69 |
| pre M - post M | 163.00 | 7.13 | 1852.83 | 22.85 | 1.83E-11 |

**Table S3.** Repeatability of connection strengths; indicative of the data presented in Figure S3. Lower and Upper represent the 95% credibility intervals of the repeatability estimate; “units” refers to the residual variance.

| Model | Component | Mode | Lower | Upper |
| --- | --- | --- | --- | --- |
| ID Overall | ID | 0.159 | 0.091 | 0.205 |
| ID Overall | units | 0.841 | 0.795 | 0.909 |
| ID:Hurricane | ID | 0.002 | 0 | 0.032 |
| ID:Hurricane | ID:PostMaria | 0.76 | 0.721 | 0.791 |
| ID:Hurricane | units | 0.233 | 0.202 | 0.26 |
| ID Before | ID | 0.408 | 0.348 | 0.478 |
| ID Before | units | 0.592 | 0.522 | 0.652 |
| ID After | ID | 0.529 | 0.367 | 0.668 |
| ID After | units | 0.471 | 0.332 | 0.633 |

